# NETosis and Myeloperoxidase Promotes Inflammation and Cardiac Remodeling in Arrhythmogenic Cardiomyopathy

**DOI:** 10.64898/2026.04.14.718596

**Authors:** Emily A. Shiel, Gallage H D Nipun Ariyaratne, Waleed Farra, Andrea Villatore, Elisa Cannon, Stephen P. Chelko

## Abstract

**Background:** Arrhythmogenic cardiomyopathy (ACM) is a heritable nonischemic cardiomyopathy and a leading cause of sudden cardiac death. Although inflammation is a pathological hallmark of ACM, the contribution of peptidylarginine deiminase 4 (PAD4)-dependent neutrophil extracellular trap (NET) formation and myeloperoxidase (MPO) to disease progression remains poorly defined.

**Methods:** To define the role of PAD4-dependent NETosis and MPO signaling in ACM disease progression homozygous desmoglein-2 mutant (*Dsg2*^mut/mut^) mice were utilized. We employed genetic and pharmacological approaches to determine the efficacy of targeting PAD4 and MPO on cardiac function, arrhythmogenic burden, myocardial fibrosis, inflammatory signaling, and gap junction integrity. Cardiac phenotyping included echocardiography, electrocardiography, histology, inflammatory profiling, and biochemical assays.

**Results:** Markers of PAD4-dependent NETosis were elevated in *Dsg2*^mut/mut^ hearts as early as 4 weeks of age, prior to cardiac dysfunction. Genetic deletion of *Pad4* significantly preserved left ventricular function, reduced ectopics, attenuated myocardial fibrosis, and suppressed proinflammatory and profibrotic cytokines. MPO levels were increased in *Dsg2*^mut/mut^ hearts, and genetic ablation of *Mpo* preserved cardiac function, reduced arrhythmic burden, prevented myocardial fibrosis, and restored connexin-43 phosphorylation and localization. Furthermore, pharmacological MPO-inhibition improved cardiac function, reduced arrhythmias, and attenuated inflammatory signaling, though myocardial fibrosis was not fully prevented. Notably, hearts from patients with ACM demonstrated increased MPO signal in both cardiomyocytes and non-cardiomyocyte populations compared with donor controls.

**Conclusions:** PAD4-dependent NETosis and MPO signaling are key drivers of inflammation, fibrosis, and arrhythmogenesis in early disease onset in ACM. Targeting neutrophil-mediated pathways represents a promising therapeutic strategy to mitigate disease progression in ACM.

**Clinical Perspective:** *What Is New?:*

- PAD4-dependent NET formation is activated early in ACM and directly contributes to myocardial inflammation, fibrosis, arrhythmias, and cardiac dysfunction.
- Genetic ablation of *Pad4* or *Mpo* preserves cardiac function, reduces arrhythmogenic burden, and attenuates proinflammatory and profibrotic signaling in a *Dsg2* mutant model of ACM.
- Pharmacological inhibition of MPO improves cardiac function and electrical stability, identifying neutrophil-derived pathways as modifiable drivers of disease.

*What Are the Clinical Implications?:*

- Neutrophil-mediated inflammation represents a clinically relevant mechanism in ACM that may be targeted without global immunosuppression.
- MPO inhibition may offer a novel disease-modifying strategy to reduce arrhythmias and preserve cardiac function in patients with ACM.
- Neutrophil- and NET-associated biomarkers may improve early risk stratification and therapeutic decision-making in genetically susceptible individuals.

**Graphical Abstract:** 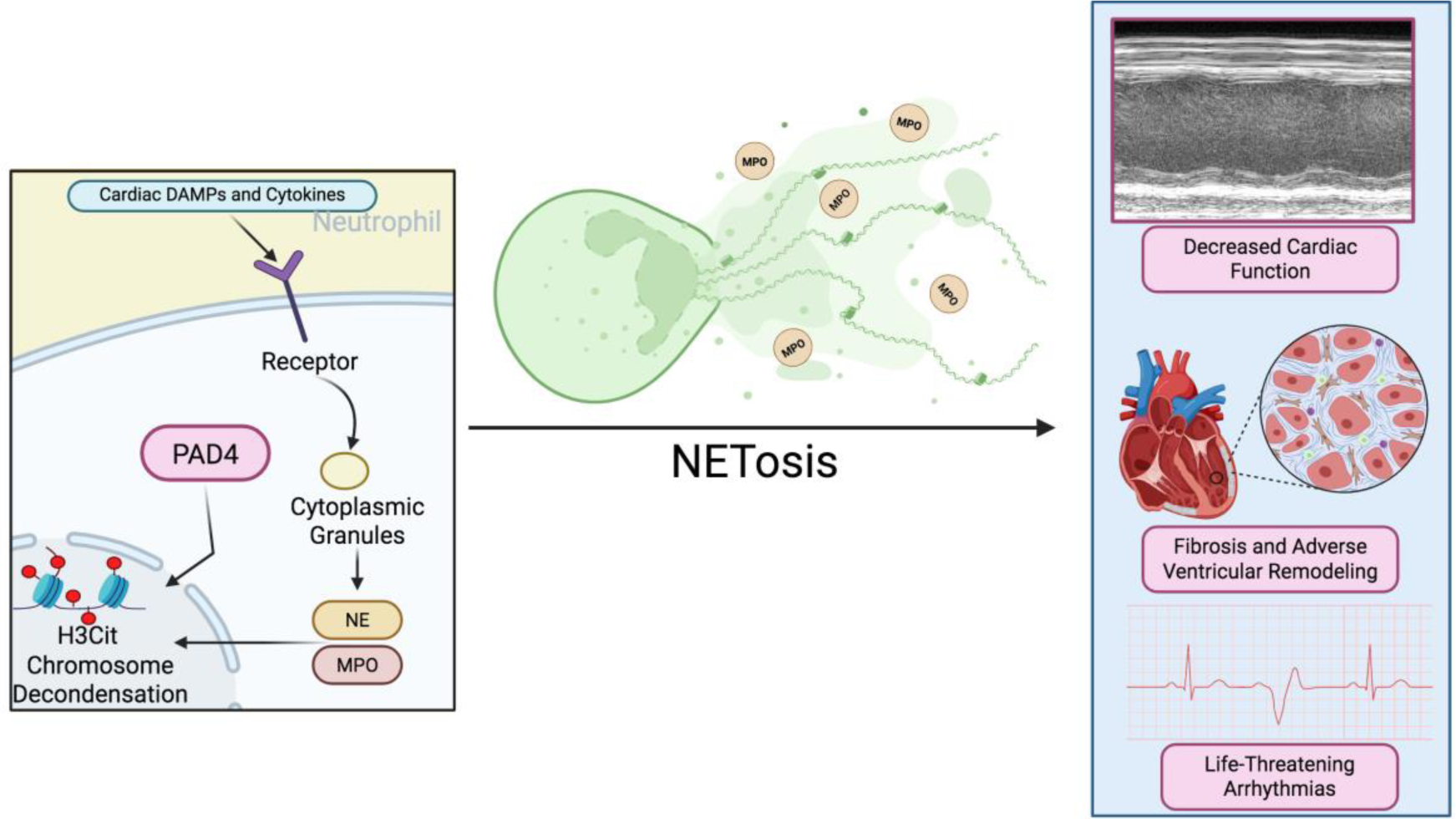

(A) Signaling pathway for PAD4-dependent NETosis. (B) Illustration of neutrophil undergoing NETosis resulting in the release of MPO and DNA histone complexes. (C) Effects of MPO release on cardiac tissue of ACM mice

## Introduction

Arrhythmogenic cardiomyopathy (ACM) is a genetic, nonischemic heart disease and is one of the leading causes for sudden cardiac death (SCD) in the young and in athletes.^1^ With a prevalence from 1:1,000 to 1:5,000, patients with ACM present with clinical symptoms such as exertional syncope, arrhythmias, chest pain, cardiac dysfunction, and heart failure (HF); and pathological features that include myocardial fibrosis and inflammation, cardiomyocyte loss, and adipogenesis.^2^ Over fifty percent of patient cases arise from desmosomal gene variants, where plakophilin-2 (*PKP2*) and desmoglein-2 (*DSG2*) are the first and second most prevalent desmosomal genes associated with ACM, respectively.^3,4^ That said, *DSG2* variants result in a more severe form of ACM and an increased risk of HF.^4^ The cardiac desmosome is a cell-cell junctional complex that is primarily responsible for maintaining mechanical and electrical continuity between cardiomyocytes.^5^

Myocardial inflammation has long been recognized as a pathological hallmark of ACM and is generally believed to be the result of infiltrating immune cells.^6,7^ Considering inflammatory infiltrates are observed in >70% of patients with ACM, this dreadful disease is frequently misdiagnosed as myocarditis.^8–10^ Prior studies demonstrated that inflammatory signaling is upregulated in ACM and is a key driver of disease progression.^11^ More specifically, research has shown elevated levels of circulating inflammatory cytokines in patients with ACM, and cardiomyocytes (CM) harboring ACM disease alleles produce and secrete potent chemotactic molecules in iPSC-CMs.^11–14^ This suggests that myocardial inflammation in ACM is mediated by both infiltrating immune cells and an innate immune response originating from CMs.

These early studies linked desmosomal disruption with suppression of the canonical Wnt/β-catenin pathway and GSK3β activation,^15,16^ and subsequent overactivation of the NFκB pathway.^11^ Outcomes from prior works demonstrated myocardial inflammation in ACM is facilitated by NFκB-induced transcription of potent inflammatory cytokines and chemokines, which promotes local inflammation and drives chemotaxis of immune cells to the heart.^11,13,14^ In fact, therapeutic blockade of NFκB was able to preserve cardiac function, prevent cardiac fibrosis and inflammation, temper myocardial levels of potent inflammatory mediators, and blunt cardiomyocyte cell death.^11^ However, long-term NFκB inhibition would likely result in severe and systemic immunosuppression, making it an unlikely target for therapeutic intervention. Thus, the field has pivoted to downstream targets such as chemotactic mediators that influence macrophage infiltration, local, cardiomyocyte specific signaling, and/or blocking NFκB-mediated transcripts as potential therapeutic targets.

One such study discovered that regulatory T cells from patients with ACM produce IL-32 which instigated lipid droplet accumulation and collagen deposition *in vitro*, suggesting T cells may also be a contributing immune cell to ACM disease progression.^17^ Conversely, preventing CCR2+ macrophage infiltration via germline deletion of *Ccr2* in a *Dsg2* mutant model of ACM showed promise in the prevention of ACM disease progression.^13^ This study first demonstrated that immune cell populations such as CCR2+ macrophages migrate to ACM hearts at various time points, indicating a diverse immune response throughout the duration of disease.^13^ While this study showed marked reduction in myocardial fibrosis and arrhythmias, *Ccr2* knockout failed to preserve cardiac function; suggesting other immune cell populations and their respective signaling pathways may be contributing factors to overall ACM disease onset and progression.^13^

In recent years neutrophils have been shown to play a major role in both ischemic and nonischemic heart diseases.^18–22^ Particularly, many of these studies highlight the role of neutrophil extracellular trap (NET) formation as a major regulator of disease progression. For example, studies using both patient data and rodent models of myocarditis, myocardial infarction, atherosclerosis, and heart failure have shown that NETosis-regulated signaling cascades are elevated and correlated with a more severe disease progression.^22–25^ NETosis can result in neutrophil cell death or vital NET release, and there are many mechanisms for NET formation that are currently known. However, a variety of studies suggest that the peptidylarginine deiminase 4 (PAD4)-dependent mechanism of NETosis plays a significant role in cardiovascular diseases (CVDs); therefore, this study aimed to unravel this mechanism in ACM. PAD4-dependent NET formation is induced by elevated intracellular calcium levels.^26^ Such that when intracellular calcium levels are high, PAD4, a calcium-dependent enzyme, is activated.^26^ Once activated, PAD4 translocates from the cytosol into the nucleus and PAD4-mediated citrullination of histone 3 (H3Cit); triggering neutrophil DNA decondensation.^27^ Myeloperoxidase (MPO) and neutrophil elastase (NE) also translocate to the nucleus to promote chromatin unfolding and nuclear membrane disruption. Following nuclear membrane disruption, the unfolded DNA is decorated with cytoplasmic proteins and released into the interstitium.^27^ This structure of scaffolded DNA covered in NE, MPO, H3Cit, and cytoplasmic proteins forms the NETs.^27^ It is important to note that NETosis has been shown to be activated by cytokines and other molecules such a tumor necrosis factor-α (TNFα), lipopolysaccharide (LPS), and hydrogen peroxide (H_2_O_2_).^26^

Previous studies showed that MPO, one of the primary proteins released during NETosis, is upregulated in various models of ACM.^11,14^ Of importance, Lubos N and colleagues demonstrated that MPO^+^ neutrophils infiltrated the heart of ACM mice as early as 19 days of age, far earlier than pro-inflammatory macrophages.^28^ MPO is one of the most abundant proteins in neutrophils and accounts for approximately 5% of their dry weight.^29^ Recently, upregulation of MPO has been linked to vascular, liver, and cardiac dysfunction.^29^ In the heart, MPO is known to upregulate matrix metalloproteinases (MMPs), such as MMP-9, leading to adverse cardiac remodeling.^29^ Furthermore, increased levels of MPO in ischemic cardiac tissue has been linked to Cx43 degradation via MMP-7 signaling, resulting in an increase in arrhythmias.^30^

Recently, the inhibition of MPO has demonstrated potential in the management of CVDs. A recent study discovered a novel irreversible MPO inhibitor, AZD4831 that is currently being used in a phase I/II HF with preserved ejection fraction (HFpEF) and mildly reduced EF clinical trial.^31,32^ Similarly, a basic science study utilizing I/R mouse model, determined that PF1355, an oral MPO inhibitor, administration prevented adverse left ventricular remodeling, attenuated ejection fraction, and reduced cardiac inflammation in I/R mice.^33^ Given the extensive evidence supporting MPO’s role in promoting arrhythmias, inducing fibrosis, and driving oxidative stress, there is a strong rationale for investigating it as a potential therapeutic target in ACM.

## Methods

### Materials and Data Availability

Data that supports the findings of this study are available from the corresponding author on reasonable request.

### Animal Study Approval

All animals were housed in a climate-controlled facility under a 12-hour light/dark cycle with *ad libitum* access to water and standard rodent chow. All procedures involving animals were conducted in accordance with the National Institutes of Health Guide for the Care and Use of Laboratory Animals (NIH Publication No. 85–23, revised 1996) and were approved by the Florida State University Animal Care and Use Committee (ACUC Approval Number: 202000052).

### Human Study Approval

Formalin-fixed, paraffin-embedded (FFPE) blocks from human samples were approved by the Johns Hopkins School of Medicine Institutional Review Board (IRB; study no. NA_00041248, PI: Hugh Calkins). All participants either provided written informed consent (ACM patient samples) or anonymized, non-consented tissues were obtained for the TMA under IRB study no. IRB00183127 (PI: Marc Halushka). Investigators were blinded to patient genotype and phenotypes throughout experiment and analyses.

### Mouse Cohorts

Wild-type (WT) controls, *Mpo*^−/−^, *Pad4*^-/-^, and *Dsg2*^mut/mut^ mice were all on a C57BL/6J genetic background. Both male and female mice were included to account for sex as a biological variable. *Dsg2*^mut/mut^ mice were generated as previously described.^15^ Both double mutant mice: (i) *Dsg2*^mut/mut^; *Pad4*^-/-^ and (ii) *Dsg2*^mut/mut^; *Mpo*^-/-^ mice were generated by breeding *Mpo*^-/-^ or *Pad4*^-/-^ strains with *Dsg2*^mut/mut^ mice in house, until double homozygosity was achieved.

### In Vivo Drug Treatment

Prior to study enrollment, baseline echocardiography was performed, and mice were randomized into experimental groups based on percent left ventricular ejection fraction (%LVEF) to minimize allocation bias. Investigators were blinded to drug treatment cohorts and genotype throughout entirety of all experiments and analyses. For the MPO inhibition study, PF1355 (MedChemExpress; Cat No. HY-100873), a selective 2-thiouracil mechanism-based MPO inhibitor, was administered to litter-matched *Dsg2*^mut/mut^ mice and WT controls via oral gavage (50 mg/kg; twice daily; from 4 to 16 weeks of age). PF1355 was dissolved in vehicle excipient containing 40mM Tris and 10% hydroxypropyl methylcellulose (HPMC; pH 10). A separate cohort of WT and *Dsg2*^mut/mut^ mice received twice daily oral gavage of equivalent volume/kg of vehicle excipient (i.e., Placebo-treated). Placebo- and PF1355-administration ranged from 100-300µL, which was dependent upon body weight throughout the course of the 12-week treatment.

### Echocardiography

Cardiac function was evaluated using the Vevo F2 imaging platform (FUJIFILM VisualSonics, Bothell, WA, USA) equipped with an ultrasound transducer (71 MHz). Mice underwent imaging under light isoflurane anesthesia (1.5–2% vaporized in 100% O_2_) delivered via nose cone. All scans were completed within 10 minutes to minimize anesthetic exposure, and heart rates were continuously monitored to remain within the physiologic range of 400–600 beats per minute. Both parasternal long-axis, B-mode and short-axis, M-mode views were captured at the level of the papillary muscles, using a sweep speed of 200 mm/s, following established protocols.^11,13,34^ A minimum of two image sets per mouse were collected and averaged in accordance with the American Society of Echocardiography guidelines for animal studies.^35^ Image processing and functional analysis were performed using the Vevo Lab software suite.

### Electrocardiogram (ECGs)

ECG recordings were collected using an iWorx IX-BIO8 eight-lead biopotential acquisition system (iWorx, Cat. No. RS-ECG8-SA). Mice were lightly anesthetized as described above and body temperature maintained at 37°C during a 10-minute continuous ECG recording session. Signal-averaged ECGs (SAECGs) were derived from filtered Lead II traces using the LabScribe ECG Analysis Module add-on software (iWorx; Cat. No. LS-ECG). The incidence of ectopic beats, including premature ventricular contractions (PVCs) and premature atrial contractions (PACs), was quantified by calculating the percentage of abnormal beats out of the total number of beats recorded: (number of ectopic events / total beats) × 100.

### Immunohistochemistry

After terminal collection of echocardiography and ECG data, mice were euthanized, and hearts were promptly harvested. Each heart was bisected longitudinally, rinsed in chilled 1X PBS to remove residual blood, and processed for downstream applications. One half of the heart was fixed overnight in 4% paraformaldehyde, while the other half was snap-frozen in liquid nitrogen. Fixed samples were processed using the Tissue-Tek VIP 6 AI Vacuum Infiltration Processor (Sakura Finetek USA, Inc.; Torrance, CA), followed by paraffin embedding with the Tissue-Tek TEC 6 Tissue Embedder. Paraffin-embedded tissues were sectioned at 5 µm thickness, and 2–3 slices were placed on positively-charged slides.

Masson’s trichrome staining was conducted following the protocol provided by the manufacturer (Sigma Aldrich, Cat. No. HT15-1KT). For immunofluorescence analysis, slides were first brought to room temperature and deparaffinized through three 5-minute xylene washes. Rehydration followed through graded ethanol (100%, 70%, and 30%) washes and a final wash in 1X PBS. Antigen retrieval was performed using a solution containing 1% Tris-HCl and 0.1% Proteinase K (ThermoFisher, Cat No. EO0491) and incubated for 30 minutes at 37°C. Sections were then blocked in a solution of 5% BSA with 0.1% Triton X-100 for one hour at room temperature.

Primary antibody incubation was performed overnight at 4°C using goat anti-MPO (Bio-Techne Cat. No. AF3667) at 1:30 dilution, mouse anti-cTnT (ThermoFisher Cat no. MA5-12960) at 1:100, and rabbit anti-Cx43 (Cell Signaling Cat. No. 3512S) at 1:100 dilution. The following day, slides were washed in 1X PBST and incubated in the dark with species-specific secondary antibodies at 1:5000 for two hours at room temperature. Slides were washed again and left to dry in the dark for 5 minutes. ProLong Gold Antifade Mountant with DAPI (ThermoFisher, Cat. No. 936931) was applied, and coverslips were mounted and cured overnight in the dark. Imaging was performed using a Keyence BZ-X710 microscope (Keyence Corporation, Osaka, Japan) with a 60X objective.

Quantification of fibrosis was performed using ImageJ (version 1.53e) by calculating the fibrotic area as a percentage of the total myocardial area. MPO-positive staining was quantified using ImageJ. Images were split into individual channels, and a consistent threshold was utilized for the MPO channel across all samples. The percentage of MPO-positive area per high-power field (HPF) was calculated. For cell-based analysis, MPO-positive cells were quantified and normalized to total nuclei (DAPI). At least 3 fields per sample were analyzed.

### MPO activity assay

All solutions from the MPO activity assay kit (Abcam; Cat No. ab105136) were equilibrated at room temperature. First, MPO assay buffer was added to each well (20µL for sera and 10µL for cardiac lysates). (i) 20mg of freshly obtained cardiac samples (which included both RV and LV) were washed in 1X PBS and homogenized in 200 µL of MPO Assay buffer II/MPO. Samples were then centrifuged at 3000 rpm for 10 minutes to remove any insoluble material, supernatant was collected and stored on ice, then 40 µL of lysate was diluted in MPO buffer. (ii) 30 µL of sera was diluted directly in MPO buffer. For all samples, a reaction mix consisting of 40 µL of MPO Assay buffer II/MPO Assay Buffer was combined with 10 µL of Hydrogen Peroxide Solution II/MPO Substrate was added to each well. Samples were mixed on an orbital shaker and incubated at room temperature for 1 hour. After 1 hour, 2 µL of Stop Solution was added to quench samples and incubated at room temperature for 10 minutes. Then 50 µL of TNB Reagent/Standard was added to each well and mixed on an orbital shaker at room temperature for 7 minutes. All samples were then immediately measured at OD412 nm on a microplate reader.

### Cytokine Arrays

Cardiac cytokine profiling was conducted using the Proteome Profiler Mouse XL Cytokine Array Kit (R&D Systems, Cat. No. ARY028) as previously described.^36^ Frozen myocardial tissue was homogenized in RIPA lysis buffer containing protease (Sigma; Cat. No. P8849-5ML) and phosphatase inhibitors (Sigma; Cat. No. P0001-5ML) (1:100). Total protein concentration was determined by bicinchoninic acid (BCA) assay. For each sample, 200µg of protein was applied to cytokine array membranes, and processed according to the manufacturer’s protocol, including incubation with chemiluminescent substrate. Signals were visualized using the Azure Biosystems 400 imaging system, and densitometric analysis was performed with Quick Spots software (Version 25.5.1.2, Ideal Eyes Systems).

### Serum

Serum was collected in BD Microtainer™ Capillary Blood Collector and BD Microgard™ Closure (Cat. No. 02-675-185) tubes and centrifuged for 10 minutes at 4°C at 3000 rpm.

### Western Immunoblots

Protein levels in each sample were first measured using a Pierce BCA Protein Assay Kit (Thermo Fisher Cat No. 23227). 30 ug of protein was loaded into each well of a 4-12% Tris-glycine SDS PAGE gel (Invitrogen Cat No. XP04205) and ran at 120V. Gels were transferred using the iBlot 2 system (Cat No. IB21001) at 20V for 7 minutes. Blots were stained with Revert520 kit (Licor Bio P/N: 926-10011) to measure total protein and then blots were blocked for 30 minutes with iBind solution (Invitrogen Cat No. SLF2020) for 30 minutes. Rabbit anti-Cx43 (Cell Signaling Cat. No. 3512S) at 1:1000, rabbit anti-phospho-RyR2 (at Serine 2808; ThermoFisher Cat No. PA5-36758) at 1:500, and rabbit anti-total-RyR2 (Thermo Fisher Cat No. PA5-104444) at 1:500 was used along with species-specific secondaries (IRDye 800CW; Licor Bio P/N 925-32211) at 1:5000. Blots were imaged using ChemiDoc MP Imaging System (BIORAD Cat No. 12003154).

### Statistical analysis

All data is presented as mean ± standard error of the mean (SEM) with sample size (n) stated within each figure legend for all experiments. Shapiro-Wilk normality test was first performed on all data sets to determine normality. Subsequently, comparison for data with one independent variable was determined by either one-way analysis of variance (ANOVA) with Tukey multiple comparisons test, Welch’s T-test, or Mann-Whitney U Test. For comparisons with two independent variables, two-way ANOVA with Tukey multiple comparisons test used. All statistical analyses were done using R 4.2.2. and the p-value of <0.05 was considered statistically significant. Furthermore, all statistical tests performed for each data set are included within each figure legend. No data was excluded from any analyses.

## Results

### Genetic ablation of *Pad4* reduces severity of key ACM phenotypes

Our prior findings demonstrated a substantially elevated neutrophil population in the hearts of *Dsg2*^mut/mut^ mice, that “waxed and waned” throughout disease development. Yet these studies solely focused on the pathological role of CCR2+ macrophages in myocardial injury in ACM. Thus, given our prior findings, we aimed to determine the relative contributions of PAD4-mediated H3Cit on NETosis in ACM. Notably, as early as 4 weeks of age, PAD4 and H3Cit, both key indicators of NET formation, were markedly upregulated in *Dsg2*^mut/mut^ mice (Fig 1A-B). We additionally found neutrophils forming active NETs, via H3Cit+ and MPO+ immunostaining, in *Dsg2*^mut/mut^ hearts in 4-week-old cardiac tissue (Figure S1).

**Figure 1.**
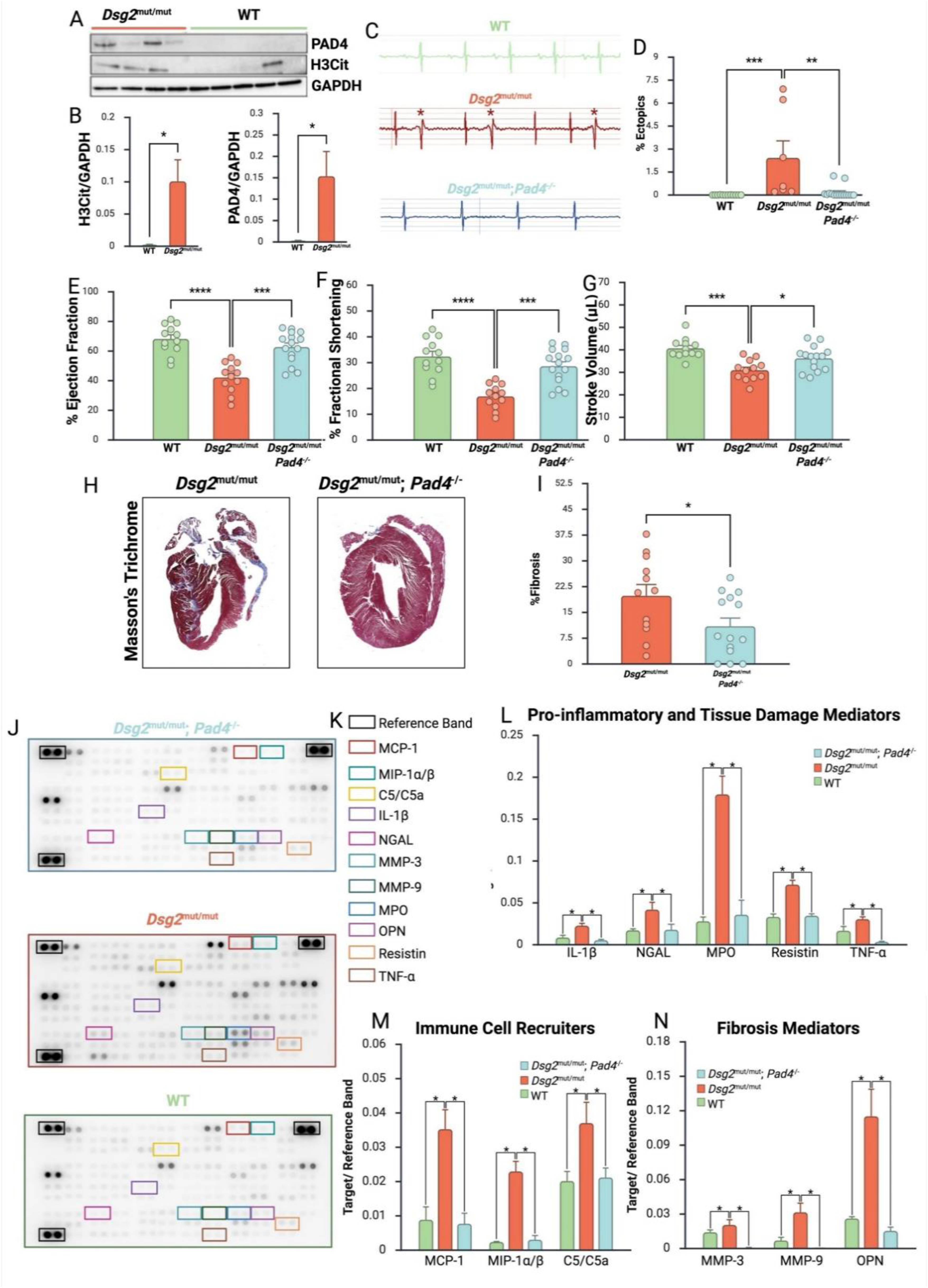
Genetic ablation of *Pad4* preserved cardiac function and reduced fibrosis at 16 weeks of age. (A) Representative Western blots clearly show the upregulation of PAD4 and H3Cit in 4-week-old *Dsg2*^mut/mut^ cardiac lysates. (B) Quantification of western blots from Fig 1A. Data presented as mean±SEM; n=4 mice/parameter; ***P<0.001 and ****P<0.0001 via two-tailed T-test. (C) Representative ECG tracings from WT, *Dsg2*^mut/mut^, and *Dsg2*^mut/mut^; *Pad4*^-/-^ at 16-weeks of age, respectively. (D) Percent ectopic beats at 16-weeks of age. Data presented as mean±SEM; n≥7. Statistical significance was determined via one-way ANOVA with Tukey’s multiple comparison test (**P<0.01 and ***P<0.001). (E) Percent left ventricular ejection fraction at 16-weeks of age. (F) Percent left ventricular fractional shortening at 16-weeks of age. (G) Stroke Volume at 16-weeks of age. (H) Representative Masson’s trichrome-immunostained hearts from *Dsg2*^mut/mut^ and *Dsg2*^mut/mut^; *Pad4*^-/-^ at 16-weeks of age. (I) Percent fibrosis at 16-weeks of age. Data presented as mean±SEM; n≥12. Statistical significance was determined via one-way ANOVA with Tukey’s multiple comparison test for E-G and via two-tailed T-test for I (*P<0.05, ***P<0.001, and ****P<0.0001). (J) Representative cytokine arrays with legend for key cytokines from WT, *Dsg2*^mut/mut^, and *Dsg2*^mut/mut^; *Pad4*^-/-^ mice. (K) Legend of selected cytokines. (L-N) Quantification of selected cytokines normalized to reference band in WT, *Dsg2*^mut/mut^, and *Dsg2*^mut/mut^; *Pad4*^-/-^ mice. Data presented as mean±SEM; n=4. *P<0.05, via two-tailed T-test. C5/C5a, Complement Component C5/C5a; MCP-1, Monocyte Chemoattractant Protein-1; MIP-α/β, Macrophage Inflammatory Protein-1 alpha/beta; MMP-3, Matrix metalloproteinase-3; MMP-9, Matrix metalloproteinase-9; MPO, Myeloperoxidase; NGAL, Neutrophil gelatinase-associated lipocalin; TNFα, Tumor Necrosis Factor-alpha; and OPN, Osteopontin.

Given these findings, we sought to determine whether PAD4-dependent NETosis contributes to ACM disease in 16-week-old *Dsg2*^mut/mut^ mice, an age when these mice demonstrate cardiac dysfunction, elevated arrhythmias, and extensive biventricular fibrosis.^11,13,15,37^ Therefore, we crossed a *Pad4*-null line (*Pad4*^-/-^) with *Dsg2*^mut/mut^ mice and performed echocardiograms, ECGs, and histology at 16 weeks of age to assess key ACM disease phenotypes. We found that *Dsg2*^mut/mut^; *Pad4*^-/-^ double mutant mice had reduced percentage of ectopic beats (0.18 ± 0.41%) than *Dsg2*^mut/mut^ mice (2.4 ± 2.9%) (Fig 1C-D). Furthermore, *Dsg2*^mut/mut^; *Pad4*^-/-^ double mutant mice showed preserved percent left ventricular ejection fraction (62.7 ± 2.8 %LVEF), fractional shortening (28.6 ± 1.7 %LVFS), and stroke volume (36.2 ± 1.4μl) compared to *Dsg2*^mut/mut^ mice (42.2 ± 2.9%, 16.9 ± 1.4%, and 31.0 ± 1.3μl; respectively) (Fig 1E-G). Additionally, *Dsg2*^mut/mut^; *Pad4*^-/-^ double mutant mice displayed reduced percentage of cardiac fibrosis (10.9 ± 2.3%) compared to *Dsg2*^mut/mut^ mice (19.8 ± 3.3%) (Fig 1H-I). *Dsg2*^mut/mut^; *Pad4*^-/-^ mice also harbored substantially reduced myocardial levels of several pro-inflammatory cytokines and chemokines including interleukin-1β (IL-1β), neutrophil gelatinase-associated lipocalin (NGAL), MPO, resistin, TNFα, monocyte chemoattractant protein-1 (MCP-1), macrophage inflammatory protein-1α/β (MIP1-α/β), and complement component C5/C5a (C5/C5a). Additionally, *Dsg2*^mut/mut^; *Pad4*^-/-^ mice exhibited a reduction in fibrosis promoting cytokines including MMP-3, MMP-9, and osteopontin (OPN) (Fig 1J-N).

### Elevated MPO levels precedes cardiac dysfunction

One of the critical proteins released during NETosis is MPO.^38^ Through the generation of potent reactive oxidant species, MPO contributes not only to antimicrobial defense but also to host tissue injury when dysregulated.^39^ Importantly, MPO has emerged as a critical linker between innate immune activation and cardiovascular pathology, promoting generation of ventricular arrhythmias, lipid oxidation, and adverse tissue remodeling.^30,33,40^ Previous studies have demonstrated that MPO is upregulated in both iPSC-CM cultures and the in hearts of *Dsg*2^mut/mut^ mice at 16 and 24 weeks of age.^11,37,41^ Therefore, we sought to investigate whether elevated MPO was merely an outcome of advanced disease or a pathological driver. We demonstrated here that MPO is upregulated in both the sera and cardiac tissue of *Dsg*2^mut/mut^ mice as early as 4 weeks of age (Fig 2 A-B). Additionally, MPO localization was additionally found to fluctuate in cardiac tissue from 4 to 16 weeks of age in *Dsg*2^mut/mut^ mice (Fig 2C-D).

**Figure 2.**
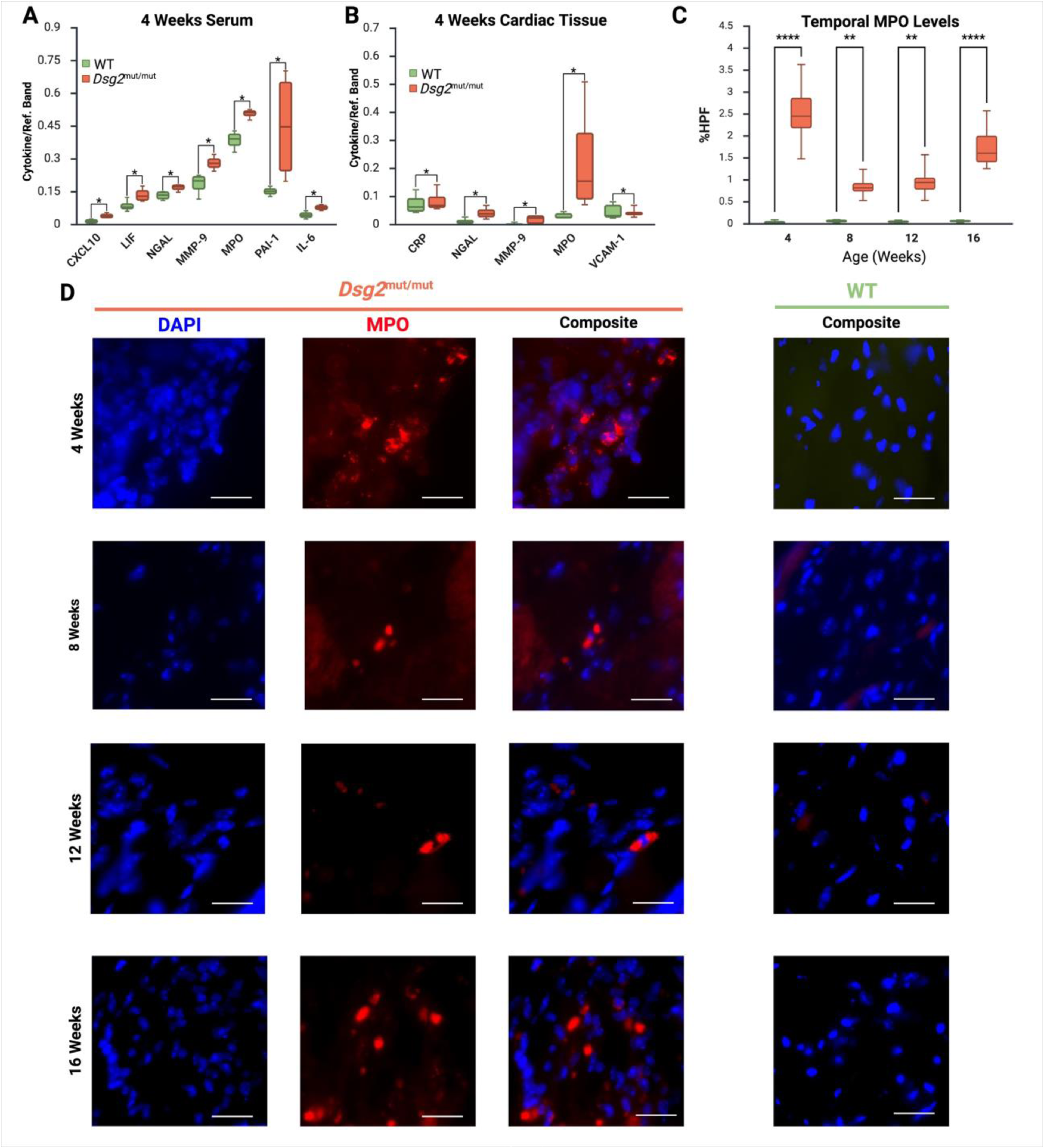
MPO is upregulated in *Dsg2*^mut/mut^ mice. Quantification of MPO levels in serum (A) and cardiac tissue (B) of *Dsg2*^mut/mut^ mice at 4-weeks of age. (C) Quantification of MPO levels in *Dsg2*^mut/mut^ cardiac tissue from part D. Data presented as mean±SEM; at least n=4 mice/parameter. Statistical significance was determined via two-tailed T-test for A-B and one-way ANOVA with Tukey’s multiple comparison test for C (*P<0.05, **P<0.01, and ****P<0.0001). (D) Representative images of hearts immunostained for MPO and DAPI from WT, *Dsg2*^mut/mut^ and *Dsg2*^mut/mut^; *Mpo*^-/-^ mice at 4-, 8-, 12-, and 16-weeks of age. Scale bar, 10µm.

### MPO-knockout attenuates ACM disease progression

To investigate the viability of targeting MPO to prevent ACM disease onset and progression we crossed *Dsg*2^mut/mut^ mice with an *Mpo*-null (*Mpo*^-/-^) line to create a double mutant mouse line (*Dsg*2^mut/mut^; *Mpo*^-/-^). These double mutants displayed a significant preservation of cardiac function at 16 weeks of age (64.8 ± 2.7 %LVEF), fractional shortening (30.0 ± 1.9 %LVFS), and stroke volume (43.9 ± 2.0μl) compared to *Dsg2*^mut/mut^ mice (42.2 ± 2.8%, 16.9 ± 1.5%, and 30.9 ± 1.3μl, respectively) (Fig 3A-C). Furthermore, *Dsg2*^mut/mut^; *Mpo*^-/-^ double mutants displayed a lower risk for arrhythmias as indicated by the significant reduction in percent ectopic beats (0.16 ± 0.1%) compared to *Dsg2*^mut/mut^ mice (2.42 ± 1.1%) (Fig 3D-E). Finally, *Dsg*2^mut/mut^; *Mpo*^-/-^ mice had significantly reduced levels of cardiac fibrosis (4.86 ± 2.0%) compared to *Dsg2*^mut/mut^ mice (19.8 ± 3.3%) (Fig 3F-G).

**Figure 3.**
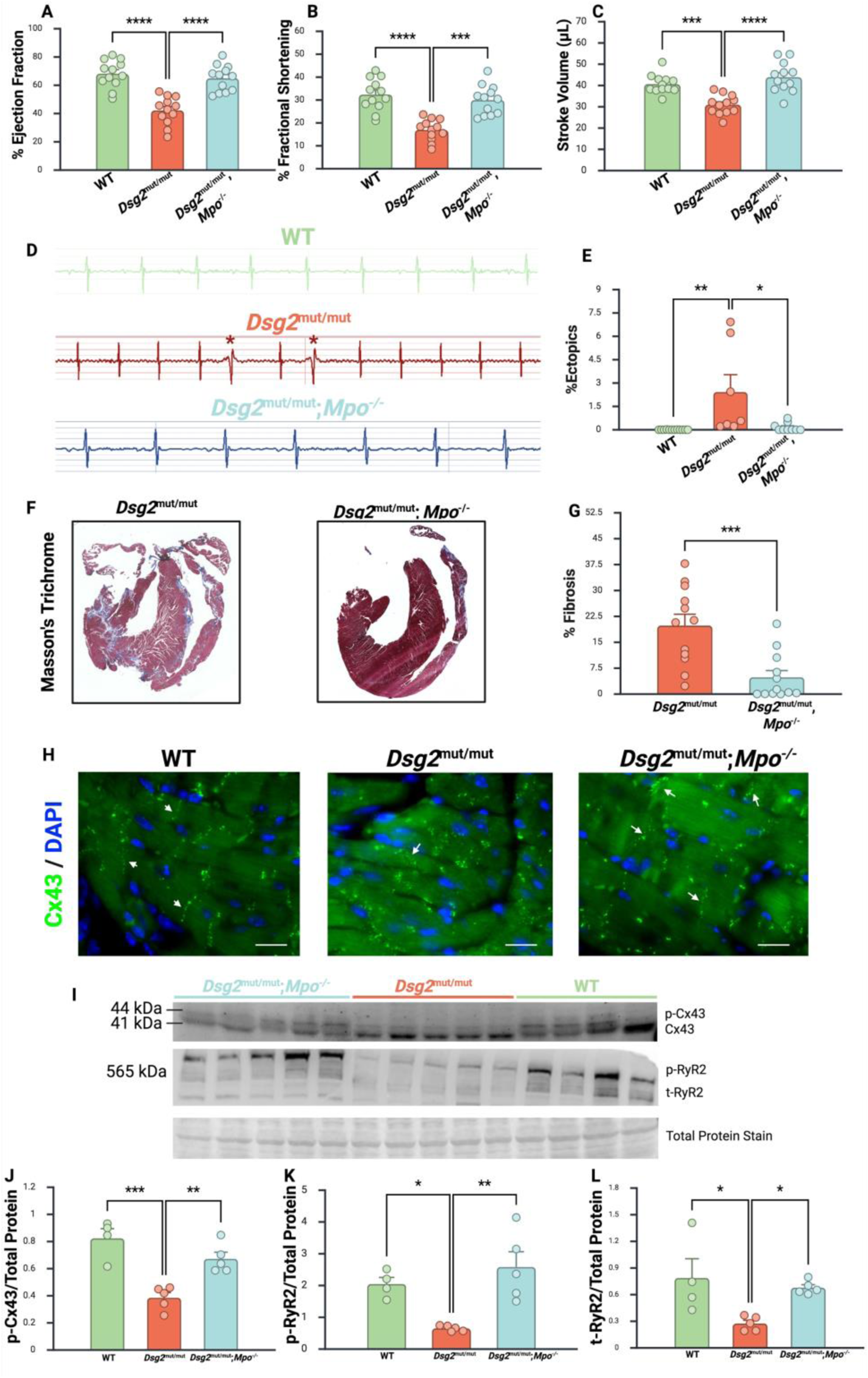
Genetic ablation of *Mpo* preserved cardiac function and reduced fibrosis at 16 weeks of age. (A) Percent left ventricular ejection fraction at 16-weeks of age. (B) Percent left ventricular fractional shortening at 16-weeks of age. (C) Stroke Volume at 16-weeks of age. (D) Representative ECG tracings from WT, *Dsg2*^mut/mut^, and *Dsg2*^mut/mut^; *Mpo*^-/-^ at 16-weeks of age, respectively. Asterisks, ectopic beats. (E) Percent ectopic beats at 16-weeks of age. Data presented as mean±SEM; n≥7. Statistical significance was determined via one-way ANOVA with Tukey’s multiple comparison test (*P<0.05, **P<0.01, ***P<0.001, and ****P<0.0001). (F) Representative Masson’s trichrome-immunostained hearts from *Dsg2*^mut/mut^ and *Dsg2*^mut/mut^; *Mpo*^-/-^ at 16-weeks of age. (G) Percent fibrosis at 16 weeks of age. Data presented as mean±SEM; n=12. Statistical significance was determined via Mann-Whitney U Test (***P<0.001). (H) Representative images of hearts immunostained for Cx43 and DAPI from WT, *Dsg2*^mut/mut^ and *Dsg2*^mut/mut^; *Mpo*^-/-^ mice. Scale bar, 20µm. (I) Representative Western blots clearly show downregulation of p-Cx43 in 16- week-old *Dsg2*^mut/mut^ cardiac lysates. (J-L) Quantification of western blot from Fig 3B. Data presented as mean±SEM; at least n=4 mice/parameter; **P<0.01 and ***P<0.001 via one-way ANOVA with Tukey’s multiple comparisons test.

A previous study has linked increased MPO activity with Cx43 degradation and disorganization in cardiovascular disease.^30^ Furthermore, Cx43 degradation and disorganization has been linked to higher risk for ventricular arrhythmias,^42,43^ slower conduction velocities,^44^ disrupted action potential propagation,^45^ and increased levels of cardiac fibrosis.^46^ Given this prior work and that mislocalization of Cx43 at the myocyte intercalated disc is a key pathological phenotype of ACM,^47^ we examined whether MPO activity impacts Cx43 levels and phosphorylation status (i.e., activity) in ACM. *Dsg*2^mut/mut^; *Mpo*^-/-^ mice displayed more organized Cx43 localization at the intercalated disc compared to *Dsg*2^mut/mut^ mice (Fig 3H). Additionally, WT and *Dsg*2^mut/mut^; *Mpo*^-/-^ mice showed increased levels of phosphorylated Cx43 than *Dsg*2^mut/mut^ mice (Fig 3I-J). Moreover, RyR2 function is known to be impaired in cardiac tissue under conditions of increased oxidative stress, a direct byproduct of MPO activity.^48^ Therefore, we assessed total and phosphorylated RyR2 (p-RyR2) levels, where *Dsg*2^mut/mut^ hearts exhibited significantly reduced phosphorylation of RyR2 at Ser2808 and total RyR2 levels compared to controls (Fig 3I and 3K-L), whereas genetic deletion of *Mpo* in *Dsg*2^mut/mut^ mice restored Ser2808 p-RyR2 and total RyR2 levels to those observed in WT hearts.

### MPO-inhibition confers cardioprotective effects in ACM

To further establish MPO as a potential therapeutic target for ACM treatment, we treated WT and *Dsg*2^mut/mut^ mice with PF1355, a selective MPO activity inhibitor (Fig 4A). Placebo-treated control mice were treated with equivalent volume/kg of vehicle. The potency of PF1355 was confirmed via MPO activity assay, which showed a 4-fold reduction in MPO activity in *Dsg*2^mut/mut^ mice (Figure S2). Of note, PF1355-treated *Dsg*2^mut/mut^ mice displayed increased ejection fraction (66.4 ± 1.6%) and fractional shortening (30.4 ± 1.0%), compared to placebo-treated mutants (56.6 ± 1.5% and 24.8 ± 1.0%; respectively) (Fig 4B-C). PF1355-treated *Dsg*2^mut/mut^ mice also showed reduced ectopics and higher R and Q amplitudes (Fig 4D-G). To determine if MPO-inhibition reduced levels of cardiac inflammatory mediators, various cytokines and chemokines were assessed. Unlike *Dsg2*^mut/mut^; *Pad4*^-/-^ and *Dsg*2^mut/mut^; *Mpo*^-/-^ double mutant mice, PF1355-treated *Dsg*2^mut/mut^ mice did not display a significant reduction in cardiac fibrosis (Fig 4H-I). *Dsg*2^mut/mut^ mice treated with PF1355 had lower levels of several pro-inflammatory cytokines and chemokines such as C5/C5a, MCP-1, MIP-1α/β, C-C motif chemokine ligand 11 and 12 (CCL11/12), IL-1β, NGAL, and lipopolysaccharide-induced CXC chemokine (LIX) (Fig 4J-M). Furthermore, levels of pro-fibrotic cytokines such as MMP-3 and MMP-9 were additionally reduced (Fig 4N).

**Figure 4.**
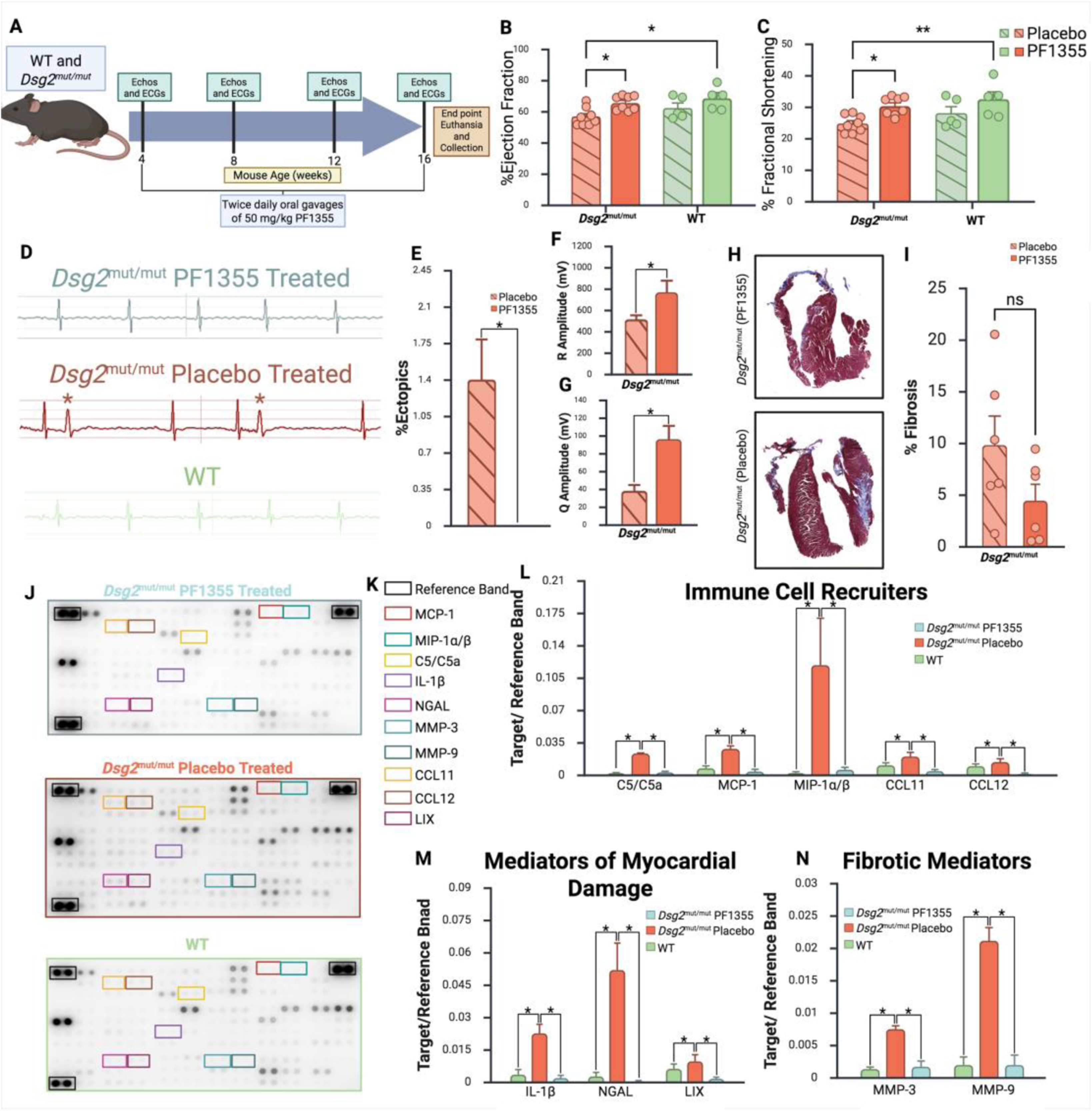
Pharmacological Inhibition of MPO activity preserved cardiac function, reduced risk of arrhythmias, and reduced inflammation at 16 weeks of age. (A) Schematic timeline of PF1355 treatment and in vivo mouse measurements. (B) Percent left ventricular ejection fraction (%LVEF) at 16-weeks of age. (C) Percent left ventricular fractional shortening (%FS) at 16-weeks of age. Data presented as mean±SEM; n≥5. Statistical significance was determined via two-way ANOVA with Tukey’s multiple comparison test (*P<0.05 and **P<0.01). (D) Representative ECG tracings from PF1355 treated *Dsg2*^mut/mut^, Placebo treated *Dsg2*^mut/mut^, and WT at 16-weeks of age, respectively. Asterisks, ectopic beats. (E) Percent ectopic beats (% Ectopics), (F) R-amplitude, and (G) Q-amplitude at 16-weeks of age. (H) Representative Masson’s trichrome-immunostained hearts from Placebo-treated and PF1355-treated *Dsg2*^mut/mut^ at 16-weeks of age. (I) Percent fibrosis at 16-weeks of age. Data presented as mean±SEM; n≥6. Statistical significance was determined via Welch’s t-test for E and one-way ANOVA with Tukey’s multiple comparison test for F,G, and I (*P<0.05). (J) Representative cytokine arrays with legend for key cytokines from Placebo-treated and PF1355-treated *Dsg2*^mut/mut^, and WT mice. (K) Legend of selected cytokines. (L-N) Quantification of selected cytokines normalized to reference band in PF1355-treated and Placebo-treated *Dsg2*^mut/mut^, and WT mice. Data presented as mean±SEM; n=4. *P<0.05, via two-tailed T-test. C5/C5a, Complement Component C5/C5a; CCL11, C-C motif chemokine ligand 11; CCL12, C-C motif chemokine ligand 12; LIX, Lipopolysaccharide-induced CXC chemokine; MCP-1, Monocyte Chemoattractant Protein-1; MIP-α/β, Macrophage Inflammatory Protein-1 alpha/beta; MMP-3, Matrix metalloproteinase-3; MMP-9, Matrix metalloproteinase-9; and NGAL, Neutrophil gelatinase-associated lipocalin. All data presented as mean±SEM. Statistical significance was determined via two-way ANOVA with Tukey’s multiple comparison test for B-C and via two-tailed T-test for E-G, I, and L-N (*P<0.05).

### Increased MPO-positive cells in hearts of patients with ACM

Prior studies demonstrated that cardiomyocytes derived from patient iPSCs harboring either a pathogenic variant in *DSG2* (∼2-fold) or *PKP2* (>3-fold) harbored elevated levels of MPO than control iPSC-CMs.^11,14^ Cardiac tissue from patients with ACM were immunostained for MPO, which displayed higher levels of cardiomyocyte MPO-positive cells (423.4 ± 10.5 MPO^+^ cells/mm^2^) compared to cardiac tissue obtained from individuals who died of non-cardiac events (38.4 ± 5.4 MPO^+^ cells/mm^2^) (Fig 5A-B). Furthermore, cardiac samples showed elevated levels of non-myocyte MPO+ cells (283.0 ± 34.7 MPO^+^ cells/mm^2^) than donor controls (20.0 ± 2.5 MPO^+^ cells/mm^2^) (Fig 5A-C). Interestingly, we observed non-myocyte MPO+ cells in samples from patients with ACM, which exhibited NET-like structures (Fig. 5C), suggestive of active NETosis.

**Figure 5.**
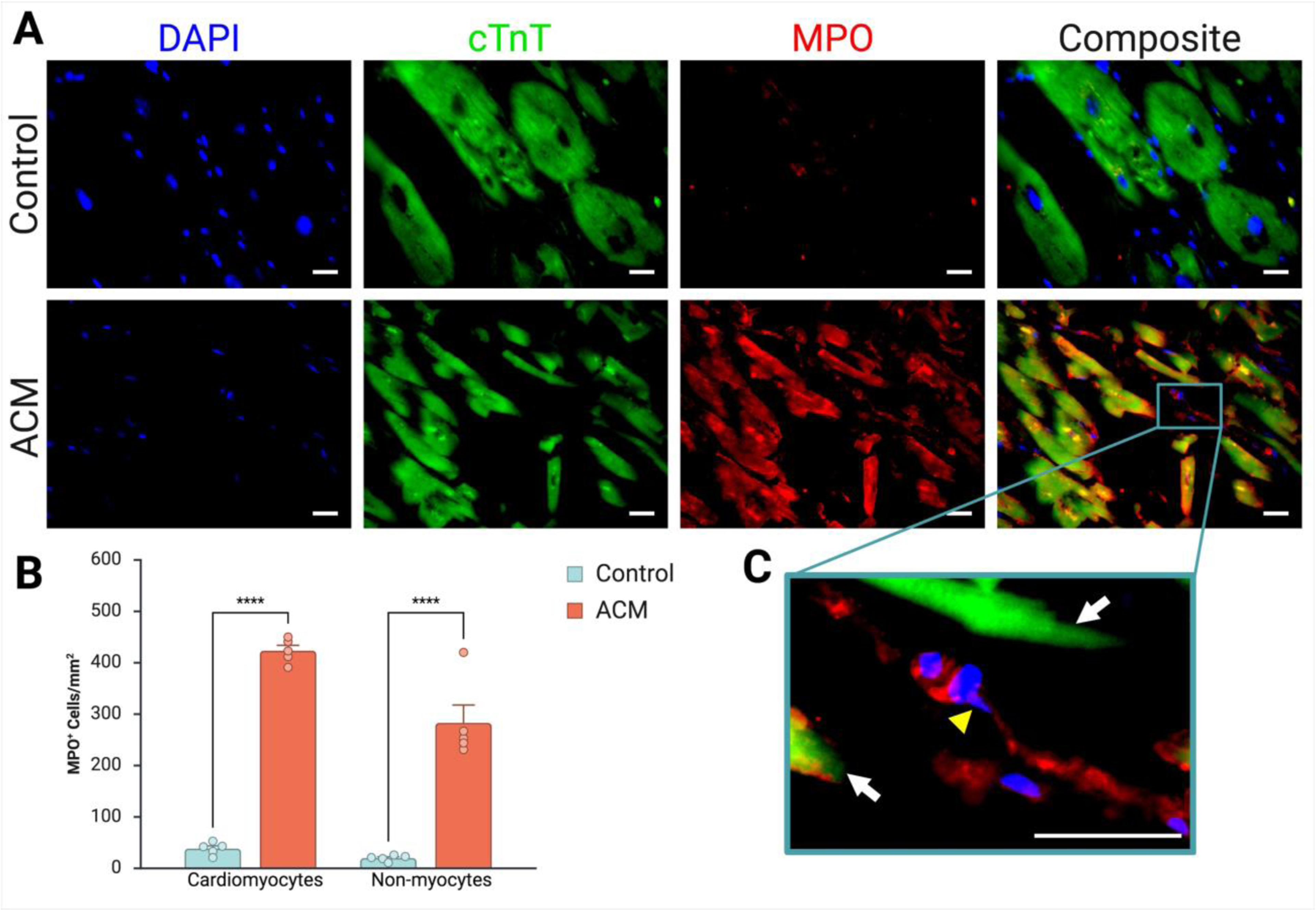
Patients with ACM have higher levels of cardiomyocyte and non-myocyte MPO. (A) Representative images of patient cardiac samples immunostained for cTnT, MPO, and DAPI from control and ACM patient cardiac biopsies. Scale bar, 20µm. (B) Quantification of MPO^+^ cells/mm^2^ from images in A. Data presented as mean±SEM; n=5; ****P<0.0001 via two-way ANOVA with Tukey’s multiple comparisons test. (C) Zoomed-in image of non-myocyte MPO^+^ cells. White arrows, cardiomyocytes; yellow arrowhead, MPO^+^ non-myocyte with NET-like projections. Scale bar, 20µm.

## Discussion

Our findings demonstrate that PAD4-mediated NETosis indeed plays a role in ACM disease progression. Complete knockout of *Pad4* in *Dsg*2^mut/mut^ mice was able to preserve cardiac function (e.g., %LVEF, %LVFS, and stroke volume) and substantially reduce arrhythmias (i.e., %ectopics), and far less cardiac fibrosis at 16 weeks of age. Additionally, cytokine array analyses revealed that *Pad4* genetic ablation in *Dsg*2^mut/mut^ mice was able to blunt the levels of key chemotactic molecules such as MCP-1, MIP-1α/β, and C5/C5a. Additionally, pro-inflammatory cytokines and mediators of tissue damage, such as IL-1β, NGAL, MPO, resistin, and TNF-α, were reduced significantly; indicating that inhibition of NET formation via PAD4 depletion was sufficient to delay cardiac inflammation in a multifaceted manner. Notably, *Pad4* knockout in *Dsg2*^mut/mut^ mice decreased key extracellular matrix and fibrotic mediators, such as MMP-3, MMP-9, and OPN. These results support the ongoing hypothesis that immune cells such as CCR2^+^ macrophages,^13^ T-cells,^17^ and now neutrophils orchestrate a pro-inflammatory cardiac environment leading to presentation of fibrosis, cardiac dysfunction, and arrhythmias in ACM subjects.

Furthermore, our findings support the role of MPO as an important player in the development of ACM disease characteristics, which is supported by the fact that both therapeutic inhibition of MPO and genetic ablation of *Mpo* preserved cardiac function and reduced arrhythmic burden. Direct MPO inhibition, via PF1355 administration, was sufficient to reduce the levels of pro-inflammatory cytokines such as IL-1β, MCP-1, C5/C5a, and LIX. Furthermore, PF1355-treated *Dsg2*^mut/mut^ mice demonstrated reduced myocardial levels of MMP-9 and MMP-3, known contributors of extracellular matrix remodeling. However, despite these beneficial outcomes, PF1355 administration failed to significantly prevent the development of myocardial fibrosis in *Dsg2*^mut/mut^ mice. Contrary to our findings from PF1355-treated *Dsg2*^mut/mut^ mice, *Dsg2*^mut/mut^; *Mpo*^-/-^ double mutant mice exhibited a significant reduction in cardiac fibrosis (4.9 ± 2.0%) compared to *Dsg2*^mut/mut^ mice (19.8 ± 3.3%). This outcome is of monumental importance, as it suggests that drug-titration of PF1355 could achieve the same effect in preventing myocardial fibrosis as that observed via *Mpo* knockout in ACM subjects. Alternatively, this outcome could indicate the temporal aspect of targeting MPO early during disease development (i.e., drug-treatment began at 4 weeks) vs complete *Mpo*-deficiency beginning at embryogenesis. Additionally, we found that genetic ablation of *Mpo* in *Dsg2*^mut/mut^ mice prevented mislocalization of Cx43 at the myocyte-myocyte intercalated disc and restored the phosphorylation levels of Cx43 to WT levels. This is particularly interesting, given that our findings above and prior works have demonstrated that increased MMP9 levels are associated with a concomitant decrease in Cx43 expression.^49^

Furthermore, although the role of RyR2 phosphorylation in arrhythmia susceptibility remains an area of active debate, our findings demonstrate that *Dsg2*^mut/mut^ hearts exhibit reduced phosphorylation of RyR2 at Ser2808. Emerging evidence links decreased pRyR2 levels with heightened risk of ventricular arrhythmias, potentially through altered sodium current reactivation and impaired calcium handling.^50–53^ Notably, genetic deletion of *Mpo* restored pRyR2 (Ser2808) levels to those observed in WT hearts. In this context, our findings demonstrating NET formation and MPO-dependent disruption of Cx43 localization, suggest that MPO promotes arrhythmogenesis and cardiac remodeling through multiple convergent pathways. Specifically, MPO-mediated oxidative inflammation may impair both intercellular electrical coupling via Cx43 and intracellular excitation–contraction coupling through dysregulation of RyR2 signaling. Collectively, these findings support a model in which MPO activity contributes to arrhythmia susceptibility in ACM by amplifying inflammatory and oxidative stress pathways that disrupt coordinated electrical and calcium signaling in the myocardium.

Importantly, these observations are not confined to animal models of ACM. Cardiac samples from patients with ACM demonstrated increased MPO signal in both cardiomyocytes and non-myocyte populations compared with donor controls, supporting a role for MPO and NETosis in the pathophysiology of ACM. In several regions, MPO staining appeared in diffuse extracellular structures consistent with NET-like formations, indicating neutrophil activation and NET release additionally occurs in human ACM myocardium. These findings therefore raise the possibility that neutrophil-driven inflammatory mechanisms, including NETosis and MPO activity, may represent an unreported contributor of ACM disease onset and progression.

While this study primarily interrogated MPO- and PAD4-dependent NETosis, additional mechanisms of NET formation, NET-associated enzymes and cytokines, and broader neutrophil effector functions remain to be explored in the context of ACM. Nonetheless, our findings establish PAD4-dependent NETosis as a previously unrecognized mediator of ACM disease. Further, our data suggests that MPO activity may serve as a central mediator of inflammatory and arrhythmogenic remodeling, although the relative contributions of other NET-derived proteins and extracellular DNA complexes will require further investigation. Moreover, potential crosstalk between neutrophils, CCR2⁺ macrophages, and other immune cell populations was not examined here, but represents an important direction for future studies. Collectively, this work expands the current immune paradigm of ACM by implicating neutrophil-driven inflammatory pathways as key modulators of disease severity and electrical instability. These findings highlight PAD4 and MPO as promising therapeutic targets and provide a foundation for the development of immune-modulating strategies to mitigate ACM progression and arrhythmic risk.

## Acknowledgments

We would like to thank Dr. Saskia Hemmers for assisting with the genotyping protocol for the *Pad4*^-/-^ mice. We would also like to thank Emily Shiel’s graduate committee members Drs. Bryant P. Chase, Jose Pinto, Michelle Parvatiyar, Jarrod Mousa, and Yi Ren for all their advice and guidance throughout this body of work. I thank Sandra Zivkovic for introducing me to NETs and for early discussions that inspired applying this concept to ACM. The authors would like to thank BioRender for the use of their illustrational and graphing tools to generate Figures 1–5, Supplemental Figures S1 and S2, and the Graphical Abstract. Finally, we are grateful to the ARVD/C patients and families who have made this work possible.

